# Discordant association of the *CREBRF* rs373863828 minor allele with increased body mass index and protection from type 2 diabetes in Māori and Pacific (Polynesian) people living in Aotearoa New Zealand

**DOI:** 10.1101/188110

**Authors:** Mohanraj Krishnan, Tanya J Major, Ruth K Topless, Ofa Dewes, Lennex Yu, John MD Thompson, Lesley McCowan, Janak de Zoysa, Lisa K Stamp, Nicola Dalbeth, Jennie Harré Hindmarsh, Nuku Rapana, Ranjan Deka, Winston W H Eng, Daniel E Weeks, Ryan L Minster, Stephen T McGarvey, Satupa’itea Viali, Take Naseri, Muagututi’a Sefuiva Reupena, Phillip Wilcox, David Grattan, Peter R Shepherd, Andrew N Shelling, Rinki Murphy, Tony R Merriman

## Abstract

**Aim/Hypotheses:** The minor allele of *CREBRF* rs373863828 associates with increased body mass index (BMI) and reduced risk of type 2 diabetes (T2D) in the Samoan population of Samoa and American Samoa. Our aim was to test *rs373863828* for association with BMI and odds of T2D, gout and chronic kidney disease (CKD) in Māori and Pacific (Polynesian) people living in Aotearoa New Zealand in 2,286 adults.

**Methods:** Association analyses were performed by linear and logistic regression with BMI, log-transformed BMI, waist circumference, T2D, gout and CKD. Analyses were adjusted for age, sex, the first four genome-wide principal components, and (when appropriate) BMI, waist circumference and T2D.

**Results:** For the minor allele of rs373863828 the effect size for log-transformed BMI was 0.038 (95% CI [0.022-0.055], *P*=4.8x10^−6^) and for T2D was OR=0.59 (95% CI [0.47-0.73], *P*=1.9x10^−6^). There was no evidence for association of genotype with variance in BMI (*P*=0.13). Nor was there evidence for association with serum urate (β=0.012 mmol/L, *P*_c_=0.10), gout (OR=1.00, *P*=0.98) or CKD (OR=0.91, *P*=0.59).

**Conclusions/interpretation:** Our results replicated, with very similar effect sizes, association of the minor allele of rs373863828 with higher BMI but lower odds of T2D among New Zealand Polynesian adults, as in Samoan adults living in Samoa and American Samoa.

## Introduction

A missense variant (rs373863828, p.(Arg457Gln)) in *CREBRF* (encoding CREB3 regulatory factor) has been associated with body mass index (BMI) in the Samoan population residing in Samoa and American Samoa [1]. The minor allele (c.1370A p.(457Gln)) associated with a 1.36kg/m^2^ higher BMI (or approximately 4kg in body weight, assuming a height of 1.7m) per copy, with an ~1.3 fold greater risk of obesity [1, 2]. In a small sample of individuals from the Kingdom of Tonga (n=171) the minor allele associated with a greater BMI of 3.1kg/m^2^ [3]. The high minor allele frequency (MAF) of the rs373863828 missense variant among Samoans in Samoa and American Samoa (MAF = 0.26) and in the Kingdom of Tonga (MAF = 0.15) [1, 3], compared to an exceedingly rare frequency in other populations in the Genome Aggregation Database (gnomad.broadinstitute.org; MAF=5.3x10^−5^ in East Asian, 3.3x10^−5^ in South Asian, 4.0x10^−5^ in European, absent in ~12K African individuals) [4], supports the hypothesis that rs373863828 is an important risk factor for obesity unique to the Samoan and Tongan populations and possibly other Polynesian populations.

Unlike *FTO-IRX3* and many other obesity risk variants in other populations, the BMI-increasing allele of rs373863828 associates with lower odds of type 2 diabetes (T2D) (OR = 0.59-0.74 after adjustment for BMI) [1]. This is contrary to the established observational association between T2D and increased BMI. Recently, however, genetic variants have also been associated with higher BMI and lower risk of T2D, hypertension and coronary artery disease in European populations [5–7]. For example, in the UK Biobank, a genetic score of eleven such alleles, including in the *IRS1* and *PPARG* loci, was associated with higher BMI (+0.12kg/m^2^) and higher body fat percentage (+0.30%) [8]. However, for a given BMI, individuals carrying these alleles were at reduced odds for T2D (OR=0.84), hypertension (OR=0.94) and heart disease (OR=0.92) [8]. These observations suggest that some molecular mechanisms that lead to higher BMI and higher body fat percentage can have ameliorating impacts on metabolic disease. In contrast, aside from fasting glucose, the CREBRF variant does not associate with other features of insulin resistance such as hypertension, lipids or homeostatic model assessment of insulin resistance [1].

In Aotearoa New Zealand, obesity and T2D are both highly prevalent in Māori and Pacific people [9]. These conditions are also strongly associated with other prevalent metabolic-based conditions in Māori and Pacific people, specifically gout and chronic kidney disease (CKD) [10–12] as complications of T2D and hypertension. Moreover increased BMI is causal of increased urate and risk of gout [reviewed in [13]] and diabetic kidney disease is ascribed as the cause of renal failure in 69% of Polynesian people [14]. *CREBRF* rs373863828 was not previously tested for association with gout or CKD [1]. Polynesian populations include those from West Polynesia (originating from Samoa, Tonga, Niue and Tokelau) and East Polynesia (Aotearoa New Zealand Māori and Cook Island Māori). Given the presence of different pathogenic allele frequencies and different linkage disequilibrium structure between East and West Polynesian populations [15, 16], understanding the genetic variation in the *CREBRF* gene in other Polynesian population groups besides those of Samoan and Tongan ancestry may provide novel insights into its association with increasing BMI, yet apparent reduction in risk of T2D.

In the present study, we tested the association of the *CREBRF* rs373863828 variant with BMI, waist circumference, T2D, gout and CKD in people of Polynesian ancestry living in Aotearoa New Zealand. In addition, based on association of *FTO* genotype with variance in BMI in Europeans [17], we tested the association of rs373863828 with variance in log-transformed BMI to detect possible underlying genetic and/or environmental influences in phenotypic variability.

## Methods

### Study population

Individuals, primarily from the Auckland, Waikato, and Christchurch regions of Aotearoa New Zealand, not known to be first-degree relatives, and aged 16 years and older, were recruited as participants of the “Genetics of Gout, Diabetes and Kidney Disease in Aotearoa New Zealand” case-control studies [18]. A separate Māori sample set from the *rohe* (area) of the Ngāti Porou *iwi* (tribe) of the Tairāwhiti region on the East Coast of the North Island of New Zealand was also included in the Aotearoa New Zealand Māori analysis. This sample-set was recruited in collaboration with Ngati Porou Hauora (Health Service) Charitable Trust. A Pukapuka Island sample set was recruited in collaboration with the Pukapuka Community of New Zealand Inc. in Mangere, South Auckland.

Information obtained at recruitment for each study included age (years), sex, height (cm), weight (kg) and waist circumference (cm) measured by trained assessors. BMI was calculated by dividing weight by the square of height in metres. Participants were also asked the ancestry of each of their grandparents. T2D was ascertained by physician-diagnosis and/or patient reports and/or use of glucose lowering therapy. Blood samples were collected and biochemical measurements were performed at the Southern Community Laboratories (<u>www.sclabs.co.nz</u>). Estimated glomerular filtration rates (eGFR) were derived from participants’ serum creatinine, age and sex using the Chronic Kidney Disease Epidemiology Collaboration equation [19]. Stage 4 and 5 CKD was defined by eGFR <30mL/min. Obesity was defined as BMI >32 kg/m^2^ [20]. Ethical approval was given by the NZ Multi-Region Ethics Committee (MEC/05/10/130; MEC/10/09/092; MEC/11/04/036) and the Northern Y Region Health Research Ethics Committee (NPHCT study; NTY07/07/074). All participants provided written informed consent for the collection of samples and subsequent analysis.

Participants who self-reported any Polynesian ancestry amongst their grandparents were separated into sample-sets based on self-reported Pacific nation of ancestry. Those participants who also reported non-Polynesian ancestry (predominantly European or Chinese) were grouped according to their Polynesian ancestry. This resulted in seven sample-sets; Aotearoa New Zealand Māori (*n*=1,296, including 270 people from the Ngāti Porou Hauora Charitable Trust study), Cook Island Māori (*n*=205), Samoan (*n*=387), Tongan (*n*=181), Niuean (*n*=47), Pukapukan (*n*=75) and an ‘Other’ Polynesian group (*n*=271), which included individuals of Tahitian (*n*=3), Tokelauan (*n*=6) and Tuvaluan (*n*=5) ancestry, along with individuals who self-reported grandparental ancestry from more than one Pacific nation (*n*=257). Pukapuka is part of the Cook Islands situated 1,140 km north-west of Rarotonga, the main island of the Cook Islands (East Polynesia), and ~720 km north-east of Samoa (Apia), geographically locating it within West Polynesia. These analysis groups were further refined based on clustering of genome-wide principal component vectors one to four (details of calculation below), resulting in the exclusion of 182 people who clustered outside of their self-reported ancestry group. The final groups used in all analyses were; Aotearoa New Zealand Māori (*n*=1,154), Cook Island Māori (*n*=197), Samoan (*n*=378), Tongan (*n*=175), Niuean (*n*=47), Pukapukan (*n*=70) and the ‘Other’ Polynesian group (*n*=265). Baseline characteristics for the final groupings are presented in Table 1.

**Table 1:** Baseline characteristics

| Characteristics | NZ Māori | Cook Island Māori | Samoa | Tongan | Niuean | Pukapukan | ‘Other’ Polynesian |
| --- | --- | --- | --- | --- | --- | --- | --- |
| **Number** | 1,154 | 197 | 378 | 175 | 47 | 70 | 265 |
| **GENDER (*n*)** |  |  |  |  |  |  |  |
| Male (proportion) | 666 (0.577) | 117 (0.594) | 280 (0.741) | 131 (0.749) | 36 (0.766) | 33 (0.471) | 162 (0.611) |
| **AGE (*years*) (SD)** | 52.25 (14.66) | 52.16 (14.28) | 45.25 (13.50) | 43.61 (14.67) | 48.28 (12.72) | 44.14 (17.28) | 42.38 (15.41) |
| **WEIGHT (*kg*) (SD)** | 96.97 (23.93) | 100.73 (25.28) | 106.90 (23.05) | 108.06 (21.61) | 96.88 (18.78) | 95.16 (20.85) | 104.63 (28.35) |
| **HEIGHT (*cm*) (SD)** | 169.57 (9.55) | 169.03 (9.99) | 173.39 (9.84) | 174.18 (8.53) | 171.73 (8.16) | 165.56 (9.34) | 171.59 (8.30) |
| **BMI (*kg/m*<sup>2</sup>) (SD)** | 33.79 (7.96) | 35.37 (8.56) | 35.63 (7.84) | 35.55 (6.88) | 32.77 (5.09) | 34.60 (6.61) | 35.43 (8.42) |
| **WAIST CIRCUMFERENCE (*cm*) (SD)** | 108.04 (18.24) | 110.75 (15.85) | 111.80 (16.28) | 112.53 (14.72) | 106.06 (12.78) | 108.20 (16.07) | 110.06 (18.81) |
| **\*SUA (*mmol/L*) (SD)** | 0.363 (0.11) | 0.406 (0.096) | 0.395 (0.13) | 0.385 (0.13) | 0.413 (0.11) | 0.405 (0.092) | 0.387 (0.11) |
| **eGFR (*mL/min*) (SD)** | 71.23 (28.42) | 69.36 (28.70) | 72.49 (27.31) | 73.00 (31.76) | 70.94 (26.68) | 79.79 (22.23) | 76.25 (28.25) |
| **#TRIGLYCERIDES (*mmol/L*) (SD)** | 2.25 (1.53) | 2.37 (1.96) | 2.13 (1.40) | 2.21 (1.43) | 2.52 (1.92) | 2.09 (1.40) | 2.10 (1.43) |
| **#HDL (*mmol/L*) (SD)** | 1.16 (0.364) | 1.17 (0.393) | 1.10 (0.340) | 1.09 (0.373) | 1.10 (0.332) | 1.21 (0.227) | 1.13 (0.366) |
| **#LDL (*mmol/L*) (SD)** | 2.80 (1.01) | 2.88 (0.99) | 2.87 (1.05) | 2.82 (1.06) | 3.03 (1.21) | 2.71 (1.05) | 2.76 (0.93) |
| **TYPE 2 DIABETES (*n*) (proportion)** | 301 (0.273) | 63 (0.330) | 83 (0.226) | 49 (0.290) | 10 (0.213) | 16 (0.229) | 61 (0.234) |
| **GOUT (*n*) (proportion)** | 456 (0.434) | 96 (0.513) | 207 (0.570) | 92 (0.532) | 21 (0.500) | 11 (0.164) | 100 (0.407) |
| **CHRONIC KIDNEY DISEASE (*n*) (proportion)** | 115 (0.119) | 23 (0.139) | 31 (0.095) | 23 (0.146) | 4 (0.111) | 0 (0.000) | 24 (0.101) |
| **rs373863828 GENOTYPE (*n*)** |  |  |  |  |  |  |  |
| G/G (proportion) | 772 (0.674) | 127 (0.651) | 212 (0.568) | 120 (0.686) | 39 (0.830) | 42 (0.600) | 168 (0.641) |
| G/A (proportion) | 348 (0.304) | 60 (0.308) | 146 (0.391) | 47 (0.269) | 7 (0.149) | 22 (0.314) | 77 (0.294) |
| A/A (proportion) | 25 (0.022) | 8 (0.041) | 15 (0.040) | 8 (0.046) | 1 (0.021) | 6 (0.086) | 17 (0.065) |
| **HWE *p*-value** | 0.049 | 0.79 | 0.098 | 0.23 | 0.34 | 0.22 | 0.052 |
| **ALLELE (*n*)** |  |  |  |  |  |  |  |
| A (proportion) | 398 (0.174) | 76 (0.195) | 176 (0.236) | 63 (0.180) | 9 (0.096) | 34 (0.243) | 111 (0.212) |

\* Serum urate concentrations are reported from individuals not taking urate-lowering therapy. # Data were unavailable for lipid-lowering medications.

### Whole genome Illumina Infinium CoreExome

The Illumina Infinium CoreExome v24 bead chip platform was used to genotype participants for ~500,000 variants across the whole-genome. Genotyping was performed at the University of Queensland (Centre for Clinical Genomics) for the Genetics of Gout, Diabetes and Kidney Disease in Aotearoa cohorts and at AgResearch (Invermay, Dunedin) for the Ngāti Porou Hauora Charitable Trust cohort. Bead chip genotyping batches were auto-clustered using GenomeStudio v2011.1 software (Illumina, San Diego). The Illumina GenomeStudio best practice guidelines and quality control protocols of Guo *et al*. were applied [21, 22]. The genotyping batches were then merged and relevant quality control steps repeated in the full dataset.

### Determination of principal components

Whole-genome principal component analysis vectors were calculated using a subset of 2,858 ancestry informative markers (as identified by Illumina) extracted from the CoreExome whole-genome genotypes. The SmartPCA (EIGENSOFT v6.0.1) [23]) program was used, with an output of 10 eigenvectors, no outlier removal, and no population size limit. Individuals of non-Polynesian ancestry were included, and the first four vectors plotted against each other to view the clustering of ancestral groupings (Asian, European, Eastern Polynesian, and Western Polynesian). The first four vectors, that explained 97.1% of the proportion of variance with the first ten vectors, were chosen for inclusion as covariates in the linear regression to account for population stratification and cryptic relatedness. Clustering by principal component vectors is presented in Figure S1.

### CREBRF rs373863828 genotyping

It was required to directly genotype rs373863828 because this variant was not present on the CoreExome genome-wide genotyping platform and we were unable to impute the region owing to the unavailability of Māori and Pacific reference haplotypes. A custom designed TaqMan™ probe-set (Applied Biosystems, Foster City, CA) was created for rs373863828 using a Python script (snp_design; DOI:10.5281/zenodo.56250) to annotate the human genome build 37 reference sequence (ftp://ftp.ensembl.org/pub/grch37) with rs373863828 and any surrounding SNPs (obtained from the NCBI dbSNP build 147 common SNP list; ftp://ftp.ncbi.nlm.nih.gov/snp). Forward Primer: CAAGAGAGGATGCTGAGACCAT; Reverse Primer:

ACCATGATGTAAGCCATTTTTCTGATACA; Probe 1 (VIC): TGAGTGGAACCGAGATAC Probe 2 (FAM): AGTGGAACCAAGATAC. Genotyping was performed using the LightCycler™ 480 Real-Time Polymerase Chain Reaction System (Roche Applied Science, Indianapolis, IN) in 384 well plates. There was a 99% successful genotyping call rate. Genotyping of 25% of the sample set as technical replicates demonstrated 100% concordance.

### Association testing

Analyses were performed using the R statistical software within RStudio v0.99.902 (<u>https://www.rstudio.com</u>). A multivariable linear regression model was used to test for association between the rs373863828 minor allele (c.1370A p.(457Gln)) and the continuous variables log-transformed BMI (a Box-Cox normality plot for the BMI data yielded λ=-0.39), untransformed BMI, waist circumference and serum urate), with the β-coefficient representing the estimated effect of each copy of the rs373863828 minor allele. For binary outcomes (obesity, T2D, gout and CKD), a multivariable binomial logistic regression model was used in a similar manner, with the allelic odds ratio (OR) representing the estimated effect of each copy of the rs373863828 minor allele. Each Polynesian population sample-set was analysed separately, and the effects combined using an inverse-variance-weighted fixed effect meta-analysis. Heterogeneity between sample-sets was assessed during the meta-analysis using Cochran’s heterogeneity (Q) statistic with random-effects analysis used when there was evidence for heterogeneity (P<0.05). For the BMI, waist circumference and T2D association analyses *P* < 0.05 was set as the significance value, given the prior probability of detecting association [1]. For the other outcomes (n=4; serum urate, gout, CKD, variance in BMI) P<0.0125 was set as the significance value to account for the multiple testing.

Models of inheritance were investigated by formulating the genotype predictor in the linear and logistic regression models (adjusting by age, sex, principal components 1-4) in different ways: additive model (0, 1, 2) - one OR; dominant model (0, 1, 1) - one OR; or recessive model (0, 0, 1) - one OR. A model selection tool (Akaike Information Criterion [24]) was used to select the most likely model. The smaller AIC value indicates the best model, but where the difference was less than two the simplest model was chosen.

### Power

Based on estimates from the Minster *et al*. [1] study and α=0.05 the power to detect an effect size of 1.36 kg/m^2^ per minor allele was >80% in the combined Māori and Pacific Island sample-set for a minor allele frequency of 0.15 or greater (Figure S2). The power to detect a moderate protective effect for T2D (OR=0.59) of the minor allele was >90% for a minor allele frequency of 0.10 or greater (Figure S2). Power calculations for gout and CKD as outcomes with α=0.0125 (to account for the four additional outcomes tested here not previously tested by Minster et al. [1] – serum urate, gout, CKD, variance in BMI) showed that power was adequate only to detect effect sizes of OR≥1.75 for gout and CKD and ≥0.032 mmol/L for serum urate (Figure S2).

### Testing for association of rs373863828 with variance in log-transformed BMI

Association of a genetic variant with variance in phenotype can detect a locus interacting in a nonadditive way without prior knowledge of the interacting factor (environmental, intrinsic, genetic). Testing for association of rs373863828 with variance in log-transformed BMI was performed as previously described [25]. The variable used as a measure of variance was produced from residuals obtained from cohort-specific analyses regressing age, age^2^ and age-by-sex. An independent ranked inverse normal transformation of absolute residuals generated z-scores, with squared z-scores (z^2^) being the variance variable. To account for the influence of rs373863828 mean-effect on the variant the mean log-transformed BMI (per genotype) was subtracted from the log-transformed BMI of each participant and the z-scores re-calculated. Linear models associating *CREBRF* genotype with both the unadjusted and mean-effect adjusted variance z-scores were performed (Equation 1). This analysis was also done on the rs373863828 genotype data of Minster *et al* [1], with age, age^2^, age-by-sex and polity as the adjusting variables in the z-score calculation steps. The Aotearoa New Zealand sample sets along with the two sample-sets of Minster *et al*. [1] were combined by an inverse-variance weighted fixed-effect meta-analysis.

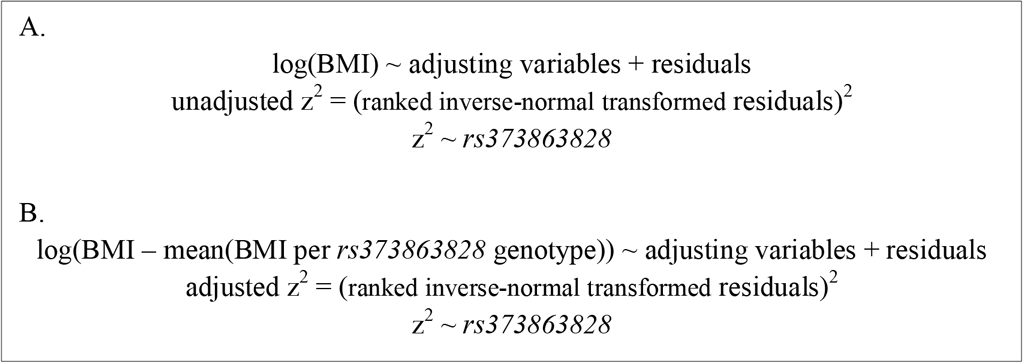

## Results

### Prevalence of rs373863828

The allele and genotype frequencies of rs373863828 in each Polynesian sample-set are presented in Table 1 and Figure 1. There was no evidence for deviation from Hardy Weinberg equilibrium in the various sample sets (Table 1; *P*>0.049). The relative frequencies of the minor (c.1370A) allele differed among the Polynesian groups, with the Samoan (MAF=0.236) and Pukapukan (MAF=0.243) groups exhibiting the highest frequency and the Niuean (MAF=0.096) group exhibiting the lowest frequency.

**Figure 1.**
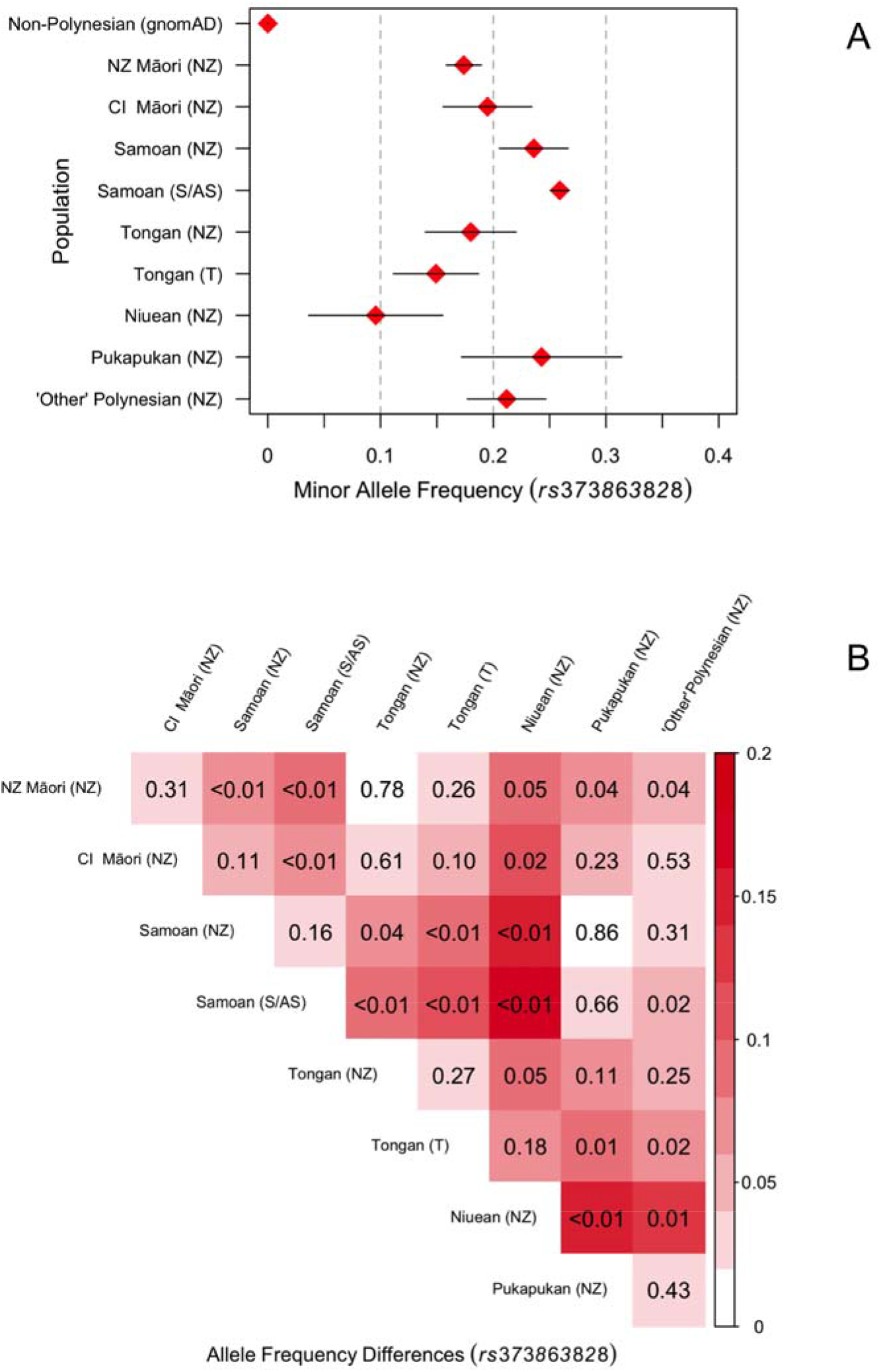
*rs373863828* allele frequencies in various Māori and Pacific ancestral groups (top) and comparison of allele frequencies between the Māori and Pacific ancestral groups (bottom). The *P* values in the bottom figure are derived from a difference in proportions parametric z-test. NZ – New Zealand. S / AS – Samoa / American Samoa (1). T – Kingdom of Tonga (3). Non-Polynesian includes all populations in the Genome Aggregation Database (gnomAD; gnomad.broadinstitute.org).

### Association analysis with adiposity measures

A fixed-effect meta-analysis of the Polynesian samples showed significant association of rs373863828 with log-transformed BMI (ß=0.038, *p*=4.8x10^−6^) with no evidence for heterogeneity between sample sets (*P*=0.19) (Tables 2 and S1; Figure 2). Association analysis with untransformed BMI revealed similar results (Tables 2, S2; Figure 2). One copy of the minor allele was sufficient to confer the effect (Table 2; β=1.80 for the heterozygote group and β=1.49 for the minor allele homozygote group compared to the major allele homozygotes in untransformed BMI analysis). This was supported by Akaike Information Criteria analysis where the dominant model had a difference of 3.3 less than the additive model. In the combined group there was association with higher odds of obesity (Table 2; OR=1.33, *p*=8x10^−4^ for >32 kg/m^2^ and OR=1.54, *p*=1x10^−5^ for >40 kg/m^2^). There was no indication of sex-specific effects (Table 2, Figure 3) and a sex-by-rs373863828 interaction analysis of the pooled sample-set showed no evidence of interaction with BMI (*P*=0.67).

**Figure 2:**
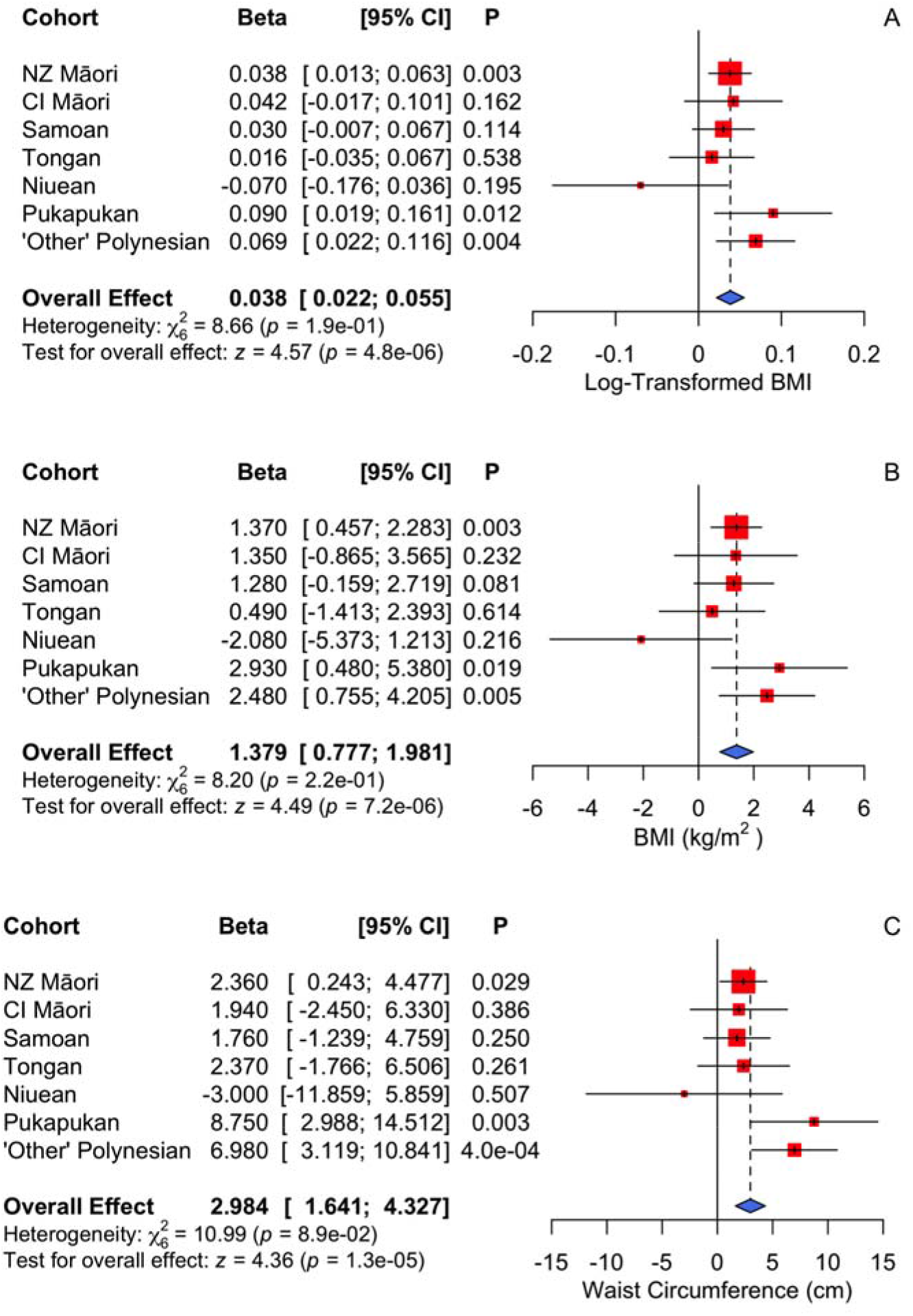
Forest plot of fixed effect meta-analysis for *rs373863828* with log-transformed BMI (A), untransformed BMI (B), and waist circumference (C). Association adjusted for age, sex, first four PCA vectors, and T2D.

**Figure 3:**
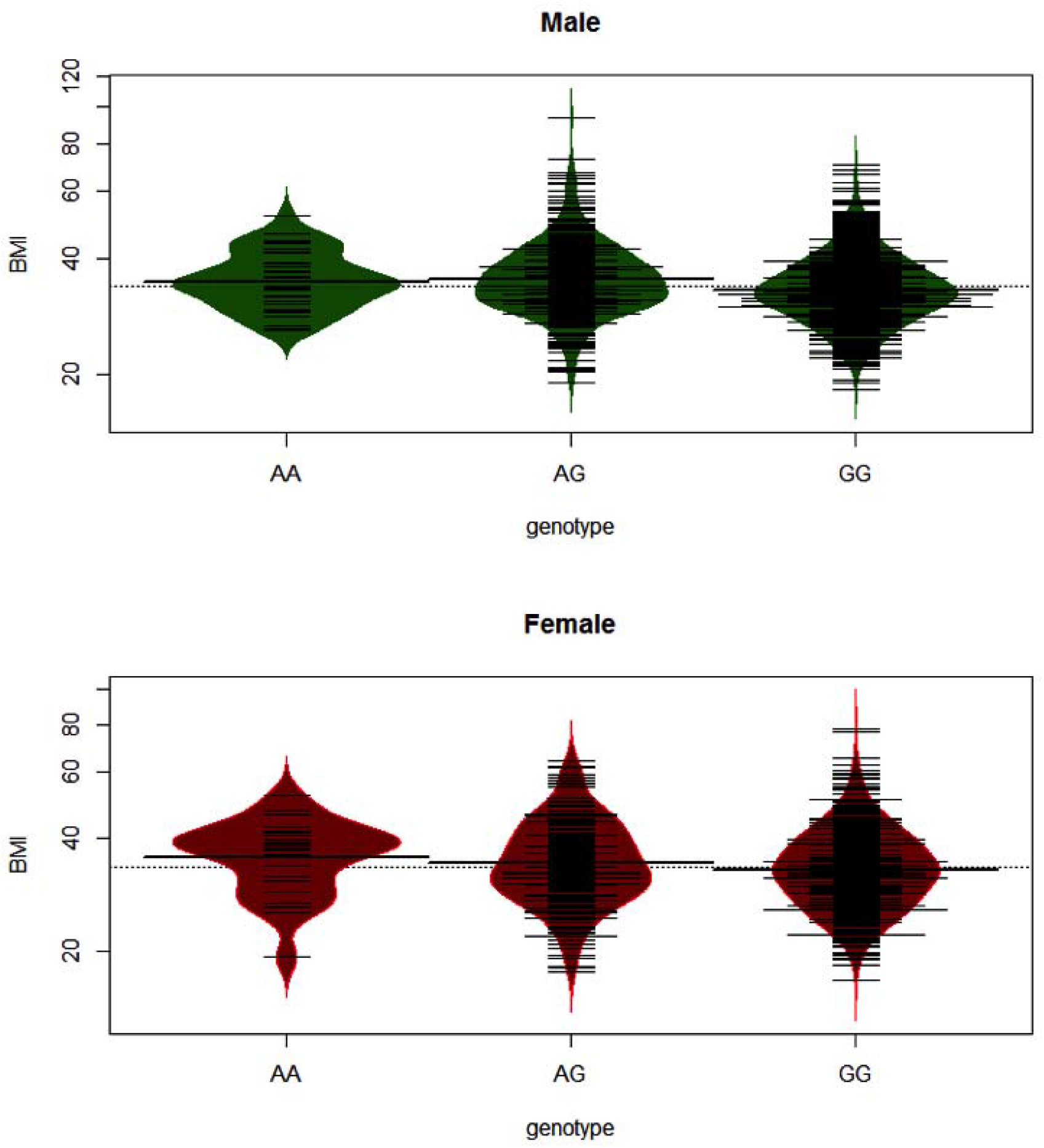
Beanplots of BMI versus *rs373863828* genotype in men and women. A solid line shows the average for each group and the dotted line the overall average. Plots were generated using the R beanplot package [29].

**Figure 4:**
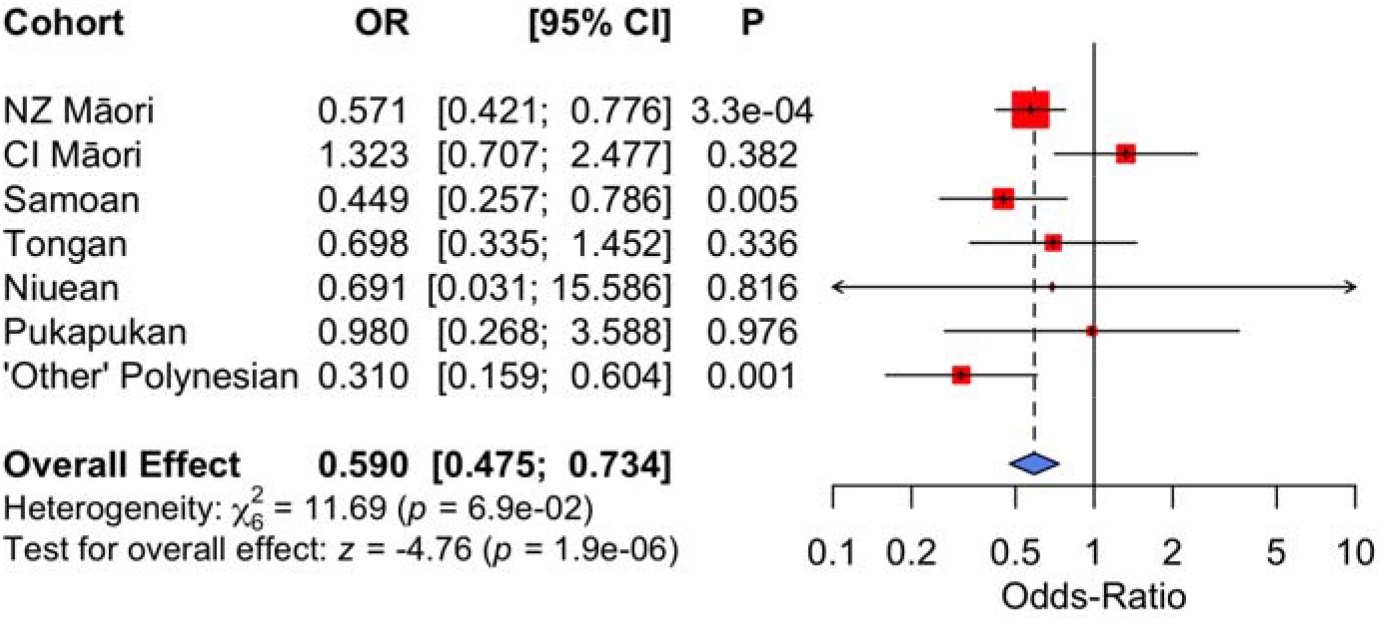
Forest plot of fixed effect meta-analysis for *rs373863828* with T2D. Association adjusted for age, sex, first four PCA vectors and T2D (OR=0.59 [0.74~0.73]), *p*=1.9x10^−6^).

**Table 2.** Rs373863828 association with weight measures, T2D, gout and CKD in the full Polynesian sample-set meta-analysis.

| Continuous trait | n | $\beta$ [95% CI] | P | Covariables |
| --- | --- | --- | --- | --- |
| Log-transformed BMI – All | 2,125 | 0.038 [0.022-0.055] | 4.8x10<sup>-6</sup> | 4 PCA, sex, age, T2D |
| Male | 1,335 | 0.042 [0.023~0.062] | 2.4x10<sup>-5</sup> | 4 PCA, age, T2D |
| Female | 790 | 0.032 [0.0014~0.062] | 0.040 | 4 PCA, age, T2D |
| Untransformed BMI (kg/m<sup>2</sup>) - All | 2,125 | 1.38 [0.78~1.98] | 7.3x10<sup>-6</sup> | 4 PCA, sex, age, T2D |
| Untransformed BMI (kg/m<sup>2</sup>) - All | 1890 | 0.16 [-0.18~0.51] | 0.35 | 4 PCA, sex, age, T2D, Waist |
| Male | 1,335 | 1.51 [0.78~2.23] | 4.4x10<sup>-5</sup> | 4 PCA, age, T2D |
| Female | 790 | 1.17 [0.089~2.26] | 0.034 | 4 PCA, age, T2D |
| Major allele homozygote | 1381 | - | - | - |
| Heterozygote | 670 | 1.80 [1.08~2.53] | 9.7x10<sup>-7</sup> | 4 PCA, sex, age, T2D |
| Minor allele homozygote | 74 | 1.49 [-0.32~3.30] | 0.11 | 4 PCA, sex, age, T2D |
| Waist circumference (cm) - All | 1,904 | 2.98 [1.64~4.33] | 1.4x10<sup>-5</sup> | 4 PCA, sex, age, T2D |
| Waist circumference (cm) - All | 1,890 | 0.66 [-0.11~1.42] | 0.092 | 4 PCA, sex, age, T2D, BMI |
| Male | 1,221 | 2.81 [1.23~4.40] | 5.0x10<sup>-4</sup> | 4 PCA, age, T2D |
| Female | 683 | 3.29 [0.80~5.78] | 9.7x10<sup>-3</sup> | 4 PCA, age, T2D |
| Urate (mmol/L)* | 1,237 | 0.015 [0.005~0.026] | 5.0x10<sup>-3</sup> | 4 PCA, sex, age |
| Urate (mmol/L)* | 1,199 | 0.012 [0.0014~0.022] | 0.026 | 4 PCA, sex, age, BMI |
| Dichotomous trait | n | OR [95% CI] | P | Covariates |
| Obesity (>32 kg/m<sup>2</sup>) | 2,125 | 1.33 [1.13~1.58] | 8x10<sup>-4</sup> | 4 PCA, sex, age, T2D |
| Obesity (>40 kg/m<sup>2</sup>) | 2,125 | 1.54 [1.27~1.87] | 1.0x10<sup>-5</sup> | 4 PCA, sex, age, T2D |
| Type 2 diabetes - All | 2,190 | 0.65 [0.53~0.80] | 3.4x10<sup>-5</sup> | 4 PCA, sex, age |
| Type 2 diabetes - All | 2,125 | 0.59 [0.47~0.73] | 1.9x10<sup>-6</sup> | 4 PCA, sex, age, BMI |
| Male | 1,335 | 0.54 [0.41~0.72] | 1.5x10<sup>-5</sup> | 4 PCA, age, BMI |
| Female | 779 | 0.66 [0.46~0.95] | 0.026 | 4 PCA, age, BMI |
| Major allele homozygote | 1,381 | - | - | - |
| Heterozygote | 670 | 0.55 [0.43~0.71] | 3.2x10<sup>-6</sup> | 4 PCA, sex, age, BMI |
| Minor allele homozygote | 74 | 0.72 [0.34~1.53] | 0.39 | 4 PCA, sex, age, BMI |
| Gout | 2,114 | 1.10 [0.91~1.32] | 0.34 | 4 PCA, sex, age |
| Gout | 2,009 | 1.00 [0.81~1.22] | 0.98 | 4 PCA, sex, age, BMI, T2D |
| Chronic kidney disease | 1,849 | 0.72 [0.53~0.97] | 0.030 | 4 PCA, sex, age |
| Chronic kidney disease | 1,795 | 0.72 [0.53~0.99] | 0.045 | 4 PCA, sex, age, BMI |
| Chronic kidney disease | 1,810 | 0.86 [0.63~1.15] | 0.36 | 4 PCA, sex, age, T2D |
| Chronic kidney disease | 1,756 | 0.91 [0.65~1.28] | 0.59 | 4 PCA, sex, age, BMI, T2D |

\*Analysis excluded individuals taking urate-lowering medication

A fixed effect meta-analysis showed evidence of association between rs373863828 and increased waist circumference in the full Polynesian sample-set (ß=2.98 cm per minor allele, *p*=1.4x10^−5^) with no evidence for heterogeneity between sample sets (*P*=0.089) (Figure 2). Adjustment of the waist circumference analysis by BMI abrogated the association with waist circumference (Table 2; lowered from 2.98 to 0.66, *p*=0.092).

### Association analysis with T2D

A fixed-effect meta-analysis of the various sample sets revealed significant association with reduced odds of T2D (OR=0.65, *p*=3.4x10^−5^) that was strengthened after adjustment by BMI (OR=0.59, *p*=1.9x10^−6^) with no evidence for heterogeneity between sample sets (*P*=0.069) (Tables 2, S4; Figure 2). Similar to the observation for BMI, one copy of the minor allele appeared to be sufficient to confer the effect (Table 2: OR=0.55 for the heterozygote group compared to the major allele homozygote group). The Akaike Information Criteria analysis showed the dominant model to be similar to the additive model (0.21 less) thus the mode of inheritance for T2D was concluded to be more consistent with an additive model.

### Association analysis of rs373863828 with serum urate, gout and CKD

We tested for association with serum urate (Table 2; Fig S3). Unadjusted these was evidence for association of the minor A-allele with higher levels (β=0.015 mmol/L, *p*=0.005, P_c_=0.020), however there was none after adjusting for BMI (β=0.012 mmol/L, *p*=0.026, P_c_=0.10). Similarly there was no evidence for association with gout (OR=1.00, *p*=0.98) (Tables 2, S5, Figure S3). There was no statistically significant (P<0.0125) evidence for association with CKD either before (OR=0.72, *p*=0.030, P_c_=0.12) or after adjustment by T2D and BMI (OR=0.91, *p*=0.59) (Tables 2, S6, Figure S3). Excluding individuals with T2D from the CKD analysis did not provide evidence for association with CKD in a single analysis of all samples pooled and adjusted for the first 4 PCA vectors, age, sex and BMI (OR=1.07 [0.65-1.69], *p*=0.79).

### Association analysis of rs373863828 with variance in phenotype

The rs373863828 variant demonstrated no effect on log-transformed BMI variance at the *CREBRF* locus for any of the New Zealand sample-sets nor the two Samoan cohorts in the study of Minster *et al*. [1]. Fixed effect meta-analysis of the New Zealand (n=2282), 1990s Samoan (n=1020) and discovery Samoan (n=1876) cohorts showed no evidence of association of rs373863828 with variance in log-transformed BMI in either the unadjusted (β=-0.053, *p*=0.15) or adjusted models (β=-0.047, *p*=0.15) (Table S7, Figure S4).

## Discussion

We have now confirmed the association of *CREBRF* variant with higher BMI (1.38 kg/m^2^ per minor allele), and higher waist circumference (2.98 cm), but lower risk of T2D (OR = 0.59) in both East Polynesian (Aotearoa New Zealand and Cook Island Māori) and non-Samoan West Polynesian adults. These results are very similar to that of Samoans living in Samoa and American Samoa, with each copy of the minor allele associated with a 1.36 kg/m^2^ increase in BMI and an OR for T2D of 0.59 [1] suggesting that the main effect of the rs373863828 minor allele is relatively impervious to environment. (Fixed effect meta-analyses that included data from the Minster et al. Samoan [1] and Naka et al. Tongan [3] studies and that indicated no evidence for heterogeneity yielded for BMI β=1.43[1.17,1.68], *p*=3.8x10^−28^ and for T2D OR=0.62[0.55,0.70], *p*=1.7x10^−14^.) Consistent also with the Samoan data [1], adjustment by BMI strengthened the association with T2D in the Aotearoa New Zealand sample set (Table 2; OR=0.65 to 0.59; Minster *et al*. discovery OR=0.64 to 0.59; Minster *et al*. replication OR=0.83 to 0.74). However, adjustment by BMI removed the association with waist circumference (Table 2), indicating that the waist circumference association was driven by overall body mass distribution rather than central adiposity.

Precisely how these genetic epidemiological findings relate to the actual CREBRF-mediated molecular pathogenesis of obesity and T2D is unclear in the absence of detailed knowledge of the molecular pathways involving CREBRF and in the absence of genetic association data with detailed body composition measures as outcome. Most population genetic variants associated with generalised obesity also associate with insulin resistance, hypertension, dyslipidemia and T2D compatible with the degree of adiposity. However, there are some genetic variants which are associated with higher BMI and percentage body fat, but lower T2D along with lower insulin resistance, hypertension, circulating triglycerides, and LDL-cholesterol. These variants are also known as ‘favourable adiposity’ variants [8] due to higher subcutaneous-to-visceral adipose tissue that suggests preferential fat storage away from visceral organs. The association of the minor allele of rs373863828 with a higher BMI but reduced odds of T2D is not entirely compatible with ‘favourable adiposity’, due to the lack of association with hypertension or lipids [1]. Consistent with the reduced odds of T2D there was weak association of the minor allele with increased insulin sensitivity by homeostatic model assessment insulin resistance in the Samoan and American Samoan populations [1] (we did not have HOMA-IR data available). Cellular bioenergetics models show that the rs373863828 minor A-allele promotes lipid and triglyceride storage at a reduced energy cost in the adipocyte suggesting that the metabolic activity of *CREBRF* in fat is important [1]. Detailed clinical studies are required to clarify whether visceral and subcutaneous body fat storage depots are altered among carriers of this variant.

It is notable that, from what is understood about the physiological role of CREBRF, there is no obvious role in regulation of appetite, which is seen at *FTO-IRX3* and other loci regulating BMI in Europeans. However, *CREBRF* is widely expressed (www.gtex.org), including throughout the brain and the *CREBRF* gene is known to regulate the CREB3/Luman protein, which is localised to the endoplasmic reticulum and plays an important role in axonal regeneration [26]. Interestingly, the CREB3/Luman protein was identified through its association with herpes simplex virus-related host cell factor 1, which has led to the hypothesis that Luman may play a role in the viral emergence from latency [27]. It will be important to explore the relationship between *CREBRF* expression in the hypothalamic nuclei and the role of CREB3/Luman in the intra–axonal translation and retrograde trafficking to promote neuronal survival in response to viral stimuli and how this may relate to BMI and T2D.

Very recently a study was published reporting association of the *CREBRF* rs373863828 minor Aallele with higher BMI in a pooled multi-ethnic sample of 4,572 New Zealand children of Māori, Pacific, European and Asian ethnicity [28]. Given that the association analysis was pooled and population-specific association analyses were not reported we are unable to directly compare our data with that study. Furthermore, we note that the A-allele was reported at a prevalence of 0.015 in European children and 0.011 in Asian children [28]. These figures are remarkable when compared to the very low prevalences in the Genome Aggregation Database (MAF=5.3x10^−5^ in East Asian, 3.3x10^−5^ in South Asian, 4.0x10^−5^ in European).

This study has confirmed in adults of Māori and Pacific (Polynesian) ancestry living in Aotearoa New Zealand that the presence of each additional minor *CREBRF* rs373863828 allele is associated with 1.38 (95%CI [0.78-1.78]) kg/m^2^ higher BMI (equivalent to 4kg/allele for an individual 1.7m in height), yet a decreased odds for T2D (OR=0.59, 95%CI [0.47-0.73])). While the prevalence of both obesity and T2D is increased amongst New Zealand Māori and those of Pacific ancestry, compared to New Zealand Europeans, our study confirms that population-specific genetic variation underpins some of the inter-individual heterogeneity observed in the discordant manifestation of obesity and T2D. This study supports the need to conduct comprehensive gene-phenotype studies in populations currently under-represented in genomic studies, in which different genetically segregating pathways linking obesity and T2D clearly exist. Such studies are not only important for these populations *per se*, but are also important in illuminating the molecular biology of the pathogenesis of metabolic disease in the wider human population and have the potential to lead to novel clinical interventions.

## Acknowledgements

The authors sincerely thank participants for generously donating their time and information to this study. The authors would like to thank Jill Drake, Jordyn de Kwant, Roddi Laurence, Christopher Franklin, Meaghan House, Nancy Aupouri, Ria Akuhata, Carol Ford and Gabrielle Sexton for recruitment.

## Funding

The Health Research Council of New Zealand (Grant #’s 08/075, 10/548, 11/1075, 14/527) and the Maurice Wilkins Centre funded the New Zealand component, and the National Institute of Health funded the Samoa and American Samoa component (Grant # R01-HL093093).

## Duality of interest

The authors declare no duality of interest relevant to the material presented herein.

## Contribution statement

MK, TJM, PRS, RM, TRM contributed to the design of the study. OD, LM, JdZ, LKS, ND, JHH, NR, RD, STM, SV contributed to data collection and RKT, LY, JMDT, WWHE, DEW, RLM, PW, DG, ANS to data analysis and interpretation. MK, RM, TRM drafted the manuscript and all other authors reviewed it. The manuscript was approved by all authors.

